# Microstructural and mechanical insight into atherosclerotic plaques– an ex vivo DTI study to better assess plaque vulnerability

**DOI:** 10.1101/2022.09.20.508689

**Authors:** B Tornifoglio, R. D. Johnston, A. J. Stone, C. Kerskens, C. Lally

## Abstract

Non-invasive microstructural characterisation has the potential to determine the stability, or lack thereof, of atherosclerotic plaques and ultimately aid in better assessing plaques’ risk to rupture. If linked with mechanical characterisation using a clinically relevant imaging technique, mechanically sensitive rupture risk indicators could be possible. This study aims to provide this link – between a clinically relevant imaging technique and mechanical characterisation within human atherosclerotic plaques. *Ex vivo* diffusion tensor imaging, mechanical testing, and histological analysis were carried out on human carotid atherosclerotic plaques. DTI-derived tractography was found to yield significant mechanical insight into the mechanical properties of more stable and more vulnerable microstructures. Coupled with insights from digital image correlation and histology, specific failure characteristics of different microstructural arrangements furthered this finding. More circumferentially uniform microstructures failed at higher stresses and strains when compared to samples which had multiple microstructures, like those seen in a plaque cap. The novel findings in this study motivate diagnostic measures which use non-invasive characterisation of the underlying microstructure of plaques to determine their vulnerability to rupture.

**Statements and Declarations:** The authors have no competing interests or declarations to declare.

**Graphical abstract:** 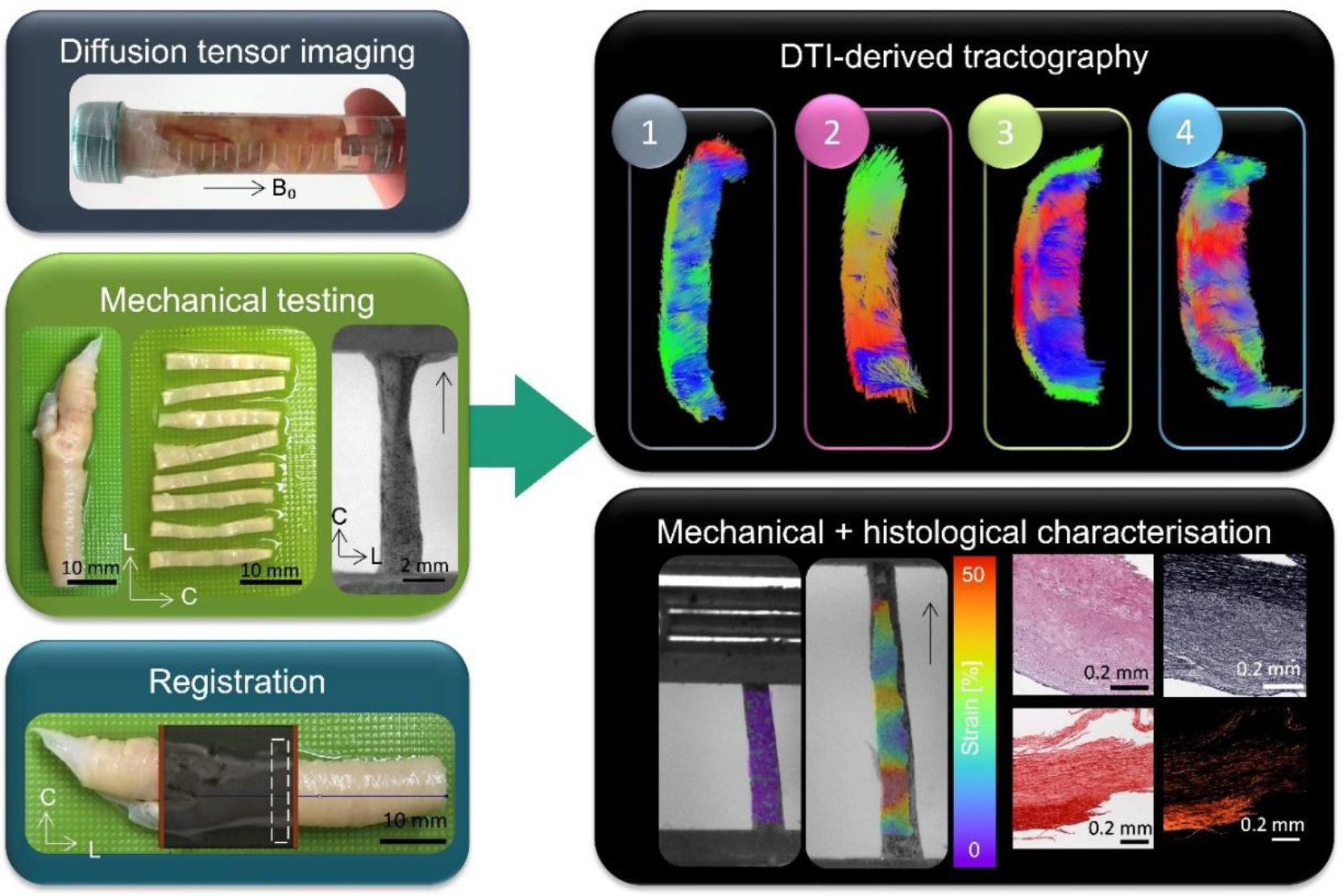

## 1.0 Introduction

There is a growing body of evidence suggesting that the percent stenosis, a diagnostic criterion for atherosclerotic plaque vulnerability established in the 1980s and 1990s^1,2^, is not an adequate indicator for measuring the likelihood of a plaque to rupture. While studies have investigated geometric features and remodeling of plaques and their usefulness in determining vulnerability^3– 6^, there is a recent acknowledgement that plaque composition and microstructure could better capitulate rupture risk^7–10^. More specifically, the percent stenosis falls short both in low grade stenosed plaques^11,12^ and asymptomatic patients^13^. A recent study identified that only 62.9% of stroke patients had carotid stenosis, and of that, 54.5% had only had mild stenosis (30 – 49%)^14^.

Initial studies investigating the composition of atherosclerosis date back to the 1960s^15^, while in the last 30 years the mechanical characterisation of plaque tissue has come to the forefront with the aim of better defining rupture risk. A few initial studies of aortic tissue highlighted the compromised mechanical integrity of calcified regions^16,17^. Calcified regions consistently fail at lower strains^16,17^. In carotid and lower limb plaques the stiffness of calcified regions has been linked to micro-computed tomography derived radiographic densities^18^. As increasing studies have been published investigating carotid, coronary, femoral and iliac atherosclerotic plaques – the highly variable nature of these tissues has become more and more apparent^17,19–22^. Due to the highly complex and dynamic microstructure within these tissues, this variability is to be expected. In the early ‘90s Lendon et al. showed that aortic plaque caps are weaker at locations with high densities of macrophages^23^, a conflicting finding to more recent work by Davis et al^24^ in carotid plaques caps. Specifically, Davis et al. showed that a higher initial stress in the plaque cap is correlated with lower macrophage content and increased collagen content. Johnston et al. showed that collagen orientation, not content, ultimately dictates the mechanical integrity of carotid plaque caps^25^. While some studies have used specific microstructural characterisation techniques, such as small angle light scattering^25^, Fourier transform infrared and scanning electron microscopy^26,27^, very few have utilised non-invasive, clinically relevant imaging^20–22^.

In comparison to ultrasound and computed tomography, magnetic resonance imaging (MRI) is both non-ionising and offers unparalleled soft tissue contrast. A number of studies have used *ex vivo* MRI to characterise plaque components^28–35^. These studies utilise different combinations of T1-, T2-, proton-density, and diffusion weighted imaging to gain insight into the plaque composition. The presence of necrotic cores^31,34^, lipid cores^28,29,31,32,35,36^, calcifications^28,30–35^, fibrous tissue^28–30,32–35^ and fibrous caps^28,31,36^, inflammation^29,31^, hemorrhage^29^, red blood cells^30^, hemosiderin^30,31^, neovascularization^31^, thrombus^32^, and solid-state and liquid-lipid^33^ have all been investigated. While knowledge of the composition of atherosclerotic plaques is beneficial in characterising them, there still lacks a direct connection between composition and mechanical integrity. As there is no current clinical imaging technique which correlates composition to mechanics, *ex vivo* mechanical characterisation of surgically excised specimens must be investigated with a imaging technique which has the potential to be applied clinically.

With previous work pointing to the significance of the alignment of load bearing collagen^25^, a non-invasive imaging technique which allows for insight into the overall microstructural alignment of a plaque could aid in characterising its integrity. Diffusion tensor imaging (DTI) is an MRI technique which characterises water diffusion within a tissue and yields insight into the microstructure. While predominantly used clinically in the brain, it has been applied *in vivo* to myocardial tissue^37,38^ and once in carotid arteries^39^. To the author’s knowledge, only one study to date by Akyildiz et al. has looked exclusively at atherosclerotic plaques ex vivo with DTI^40^. That study showed, for the first time, that a non-invasive imaging technique could characterise the overall microstructural alignment within carotid atherosclerotic plaques. However, these findings were not linked back to the mechanics of the tissue or investigated with respect to composition or specific microstructural components.

The aim of this study is to try to bridge the gap between a clinically relevant imaging technique and mechanical integrity in carotid atherosclerotic plaques. To achieve this, fresh carotid plaques from endarterectomy surgeries were imaged *ex vivo* with DTI to characterise the microstructure, then subsequently mechanically tested to failure and histologically processed. Altogether, the work presented here seeks to investigate and establish if non-invasive imaging metrics can inform the vulnerability of a plaque and ultimately surpass the percent stenosis as a clinical indicator of plaque rupture risk.

## 2.0 Methods

### 2.1 Sample acquisition

Carotid atherosclerotic plaques (n=7) were obtained from symptomatic carotid endarterectomy patients at St. James’s Hospital Dublin. All patients had a percent stenosis greater than 50%. Ethical approval was obtained from St. James Hospital ethical committee in compliance with the declaration of Helsinki. Carotid plaques were rinsed in phosphate buffered saline (PBS) to remove residual blood and cryopreserved as previously reported^25^. Samples remained at −80°C until imaging and were cryopreserved, to maintain mechanical integrity of the samples, between experimental steps.

### 2.2 Ex vivo imaging

On the day of ex vivo imaging specimens were thawed at 37°C and rinsed in PBS. Fresh plaques were secured to a custom-made 3D printed holder in a 15 ml falcon tube with fresh PBS for imaging, see Figure 1. All plaques were imaged individually and at ambient room temperature (approximately 25°C). All imaging was performed in a small-bore horizontal 7 Tesla Bruker BioSpec 70/30 USR system (Bruker Ettlinger, Germany) with a receive-only 8 channel surface coil, birdcage design transmit coil, shielded gradients (maximum strength 770 mT/m) and Paravision 6 software (Bruker, Ettlinger Germany). A conventional 3D spin echo DTI sequence, previously used on porcine carotid arteries^41^ was used. The parameters were as follows: TE/TR: 17.682/1000 ms, image size: 64 × 64 × 64, field of view: 16 × 16 × 16 mm, isotropic resolution: 250 × 250 × 250 μm, b-values: 0, 800 s/mm^2^, 10 b-directions, with fat suppression on and acquisition time: 12 hours and 30 minutes. After imaging, plaques were cryopreserved at −80°C until mechanical testing.

**Figure 1.**
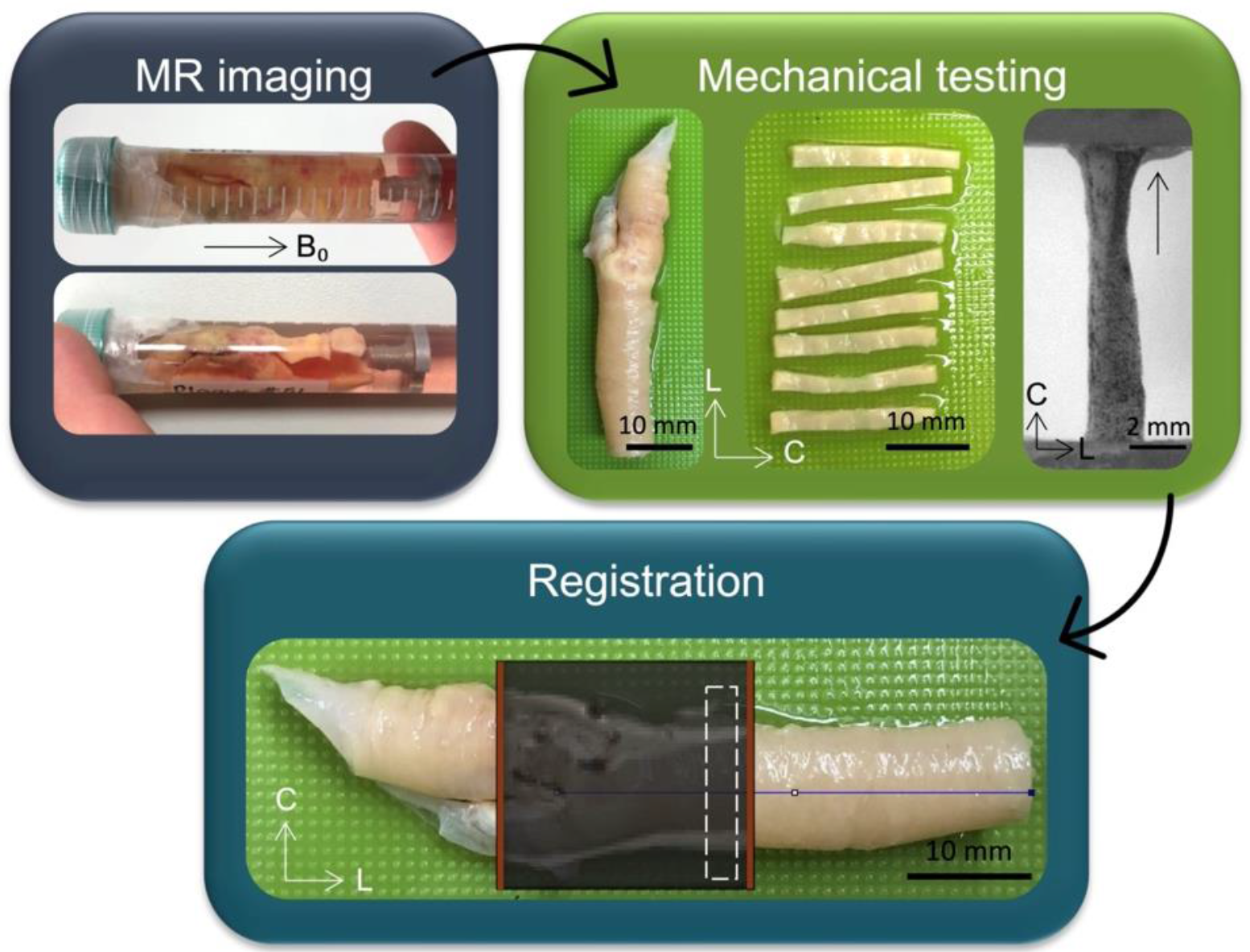
General overview of main methods in this study. Fresh carotid plaques were imaged ex vivo and then circumferential strips were sectioned and uniaxially extended to failure. MR data was then registered to specific mechanically tested strips to facilitate the investigation of DTI-metrics and mechanical properties.

### 2.3 Mechanical testing

#### 2.3.1 Circumferential strips

Samples were thawed at 37°C and rinsed in PBS. Circumferential strips (n=32) were sectioned from the plaques, each with a width of 2 mm, as seen in Figure 1. Images of each strip were taken to measure dimensions in ImageJ^42^. Three measurements were taken, and mean width and thickness were used for the calculation of the cross-sectional area. Of 32 tested strips, 15 strips were excluded from analysis due to either failure near the grips or inability to be referenced back to MR data. Of the 17 strips presented in this study, 14 strips were taken from the plaque within the common and three from plaque in the internal carotid branch. In order to use digital image correlation (DIC), strips were sprayed with a tissue marking dye (Epredia™, Fisher Scientific) using an airbrush (Kkmoon airbrush) compressor (ABEST Single Cylinder Piston Compressor).

#### 2.3.2 Uniaxial tensile tests and digital image correlation

Strips were uniaxially extended to failure using a uniaxial test machine (Zwick Z005, Zwick GmbH & Co. Ulm, Germany). All tests were performed in a PBS bath at 37°C. The testing protocol included a preload to 0.01 N, after which the force was zeroed and followed by five preconditioning cycles to 5% strain, then extension to failure. All steps were done at a strain rate of 20 mm/min. A digital image correlation (DIC) system with a two-camera set-up was used (Dantec Dynamics GmbH, Denmark) to track local strain deformations throughout the test, with camera calibration performed prior to testing. Images were acquired from both cameras at a frame rate of 5 Hz. After failure all samples were fixed in 10% formalin for histological processing.

### 2.4 Histological analysis

After fixation, strips were stepwise dehydrated (Leica TP1020, Semi-enclosed benchtop tissue processor, Germany) and embedded in paraffin wax blocks. Strips were either embedded to get 1) axial cross-sections – such that the lumen was oriented perpendicular to the face of the wax block or 2) with the luminal side of the strip flush with the face of the wax block to achieve cross-sections radially through the wall thickness. The integrity of the strip after mechanical testing dictated which of these options was most feasible; specifically, if it was not possible to orient the samples such that axial cross-sections could be obtained, the samples were oriented to get radial cross-sections. Samples were sectioned at 7 μm using Feather C35 microtome blades. Samples were stained with Haematoxylin & Eosin (H&E), picrosirius red (PSR) and Verhoeff’s elastin (Leica ST5010, Autostainer XL, Germany). Brightfield imaging of all stains was performed on an Aperio CS2 microscope with ImageScope software V12.3 (Leica Biosystems Imaging, Inc., Vista, California). Polarised light microscopy (PLM) was performed on the PSR-stained samples using an Olympus BX41 microscope with Ocular V2.0 software (Teledyne Photometrics, Tuscon, Arizona). Two PLM images were taken for each slice, 45° to each other, with the exposure time kept consistent across all samples.

### 2.5 Registration

For the co-registration, several reference points were used, namely (i) the bifurcation, (ii) the base of the plaque in the common carotid, (iii) notable calcifications, and (iv) the 3D printed holder. These reference points allowed the MR data to be segmented into individual volumes for each mechanically tested strip. Figure 1 shows an example of the MR data overlaid on the plaque, with the red lines denoting the imaging field of view. By knowing the width of the samples measured in ImageJ (2 mm approximately) and the slice thickness of the DTI images (0.25 mm), each strip corresponded to approximately eight MR slices.

### 2.6 Data processing

#### 2.6.1 DTI data analysis

All raw data was denoised and corrected for Gibbs ringing in MRtrix3 prior to fitting the mono-exponential tensor model in ExploreDTI^43,44^. From the tensor the fractional anisotropy (FA), mean diffusivity (MD), and helical angles (HA) were calculated. FA is a normalized measure of how much the eigenvalues differ and informs the degree of anisotropic diffusion within a voxel^45^. MD is the average of the eigenvalues and describes how much diffusion is occurring within a voxel. The first eigenvector (FE) was used to calculate the helical angle:

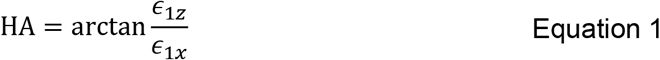

where *ϵ*_1*z*_ and *ϵ*_1*x*_ are the z- and x-components of the FE. The calculated HA represents the angle between the predominant direction of diffusion, the FE, and the plane normal to B_0_, the main magnetic field. Due to the presence of calcifications which are visualised as noise in DTI data and inherently bias the metrics^46^, some MR data was excluded from analysis. Specifically, regions of low signal (below the 50% percentile of the non-weighted image) and regions with high residuals from the tensor fitting (above the 99^th^ percentile) were removed. Lastly, the average MD of background PBS was used to remove the background and stray pixels were manually removed to yield the final tissue mask. After registration, as described in Section 2.5, 17 DTI volumes were obtained which represented the 17 mechanically tested specimens. Due to significant tissue heterogeneity, regions of the strips which were within the grips during testing were manually removed through visual inspection of images of the sample taken prior to testing and the high-resolution DIC images. Supplementary figure 1 shows the visual removal of MR data throughout these steps. At this point the final tissue regions for each mechanically tested specimen were obtained and average values of FA, MD, and HA were extracted for each specimen (one mean value and standard deviation).

#### 2.6.2 Tractography

Deterministic tractography was performed in ExploreDTI both on whole plaques as well as individual strips. For whole plaques the following parameters were used: seed point resolution: 0.25 × 0.25 × 0.25 mm, seed FA threshold: 0.075, FA tracking threshold range: 0.075-1, MD tracking threshold range: 0-infinity, linear, planar, and spherical geometry tracking threshold range: 0-1, fibre length: 2-50 mm, angular threshold: 90°, linear interpolation, and no random perturbation of seed points. For individual specimens the same parameters were refined slightly: FA threshold (0.05), FA tracking range (0.05-1), and the fibre length (1-50 mm). This refinement allowed more fibres (at a lower FA and shorter length) to be tracked. This wasn’t necessary for the larger whole plaque but gave more detail for individual strips. Specimens were visually grouped into four groups based on tractography, as follows: (1) predominantly circumferentially aligned with sparse axial tracts on the luminal side, (2) predominantly circumferential with the presence of a plaque cap shoulder, (3) circumferentially aligned medial layer with a mixed region on the luminal side, and (4) overall mixed alignment.

#### 2.6.3 Mechanical data

The cross-sectional area of each specimen was used to calculate engineering stress and the force-deformation curves after preconditioning were used to establish the stress-strain behavior of the samples. Failure was defined as the point at the first evidence of failure^47^. Specifically, when the force decreased by 5%, and the ultimate tensile (UT) stress and UT strain values were extracted. Stiffness was also calculated for each sample by taking 30 data points before the final 20% of the curve and calculating the slope of the final linear region in the stress-strain curves^48^. DIC analysis was performed on Istra4D (x64 V4.4.6.534). Evaluations were done with the following parameters: facet size: 69, 3D residuum: 10, grid spacing: 15 pixels and a low outlier tolerance. The reference frame for all DIC analysis was the frame at the end of the preconditioning cycles prior to the start of extension to failure. Engineering strain was investigated from DIC as both (1) the average strain across the gauge length on the tissue surface, called DIC strain, and (2) mean strain locally at the point of failure, called DIC local failure strain (Supplementary Figure 2).

### 2.7 Statistics

Statistical analyses were performed using GraphPad Prism (Version 8). All data was tested for normality using D’Agostino-Pearson normality tests and equality of group variances using Brown-Forsythe ANOVAs. If normality was not passed, nonparametric tests were used. Pearson’s correlations were used to determine the relationship between mechanical properties and DTI metrics, with correlation coefficients (r values) < 0.3 considered weak, 0.3 < r < 0.7 considered moderate, and r > 0.7 a strong correlation. Differences between the mechanical properties of individual subjects are included in Supplementary figure 3.

## 3.0 Results

Tractography of whole plaques can be seen in Figure 2. Red-green tracts indicate in-plane circumferential alignment, while the blue tracts represent out-of-plane axial alignment. While all specimens show some degree of axial tract alignment, there are specimens which exhibit a more disorganised alignment overall, such as the first three plaques in Figure 2. Not only is there considerable variability between different specimens, variability is also evident across individual plaques.

**Figure 2.**
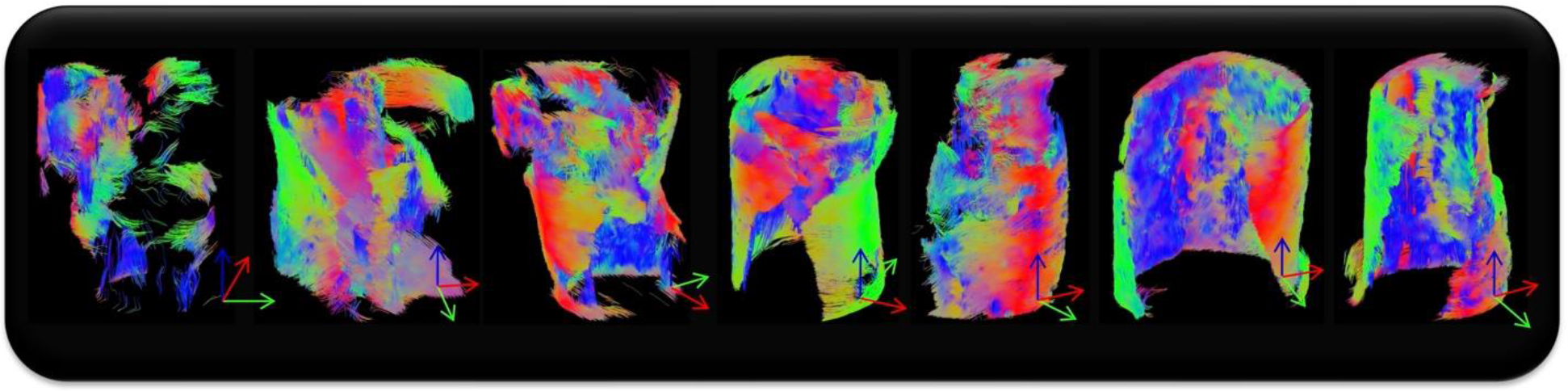
Whole plaque tractography of the seven specimens imaged and tested in this study. Red-green tracts indicate in-plane circumferential alignment whereas blue tracts represent axial out-of-plane diffusion.

Looking at the mechanical properties and DTI-derived metrics of the 17 strips, a high degree of variability was observed, see Figure 3. The mean UT stress and strain across the samples was 0.293 ± 0.2 MPa and 38.3 ± 19% strain, respectively. The mean stiffness across all samples was 1.26 ± 0.6 MPa. The mean DTI-derived MD, FA and HA were 0.0011 ± 0.0001 mm^2^/s, 0.12 ± 0.01, and 46.7 ± 6°, respectively.

**Figure 3.**
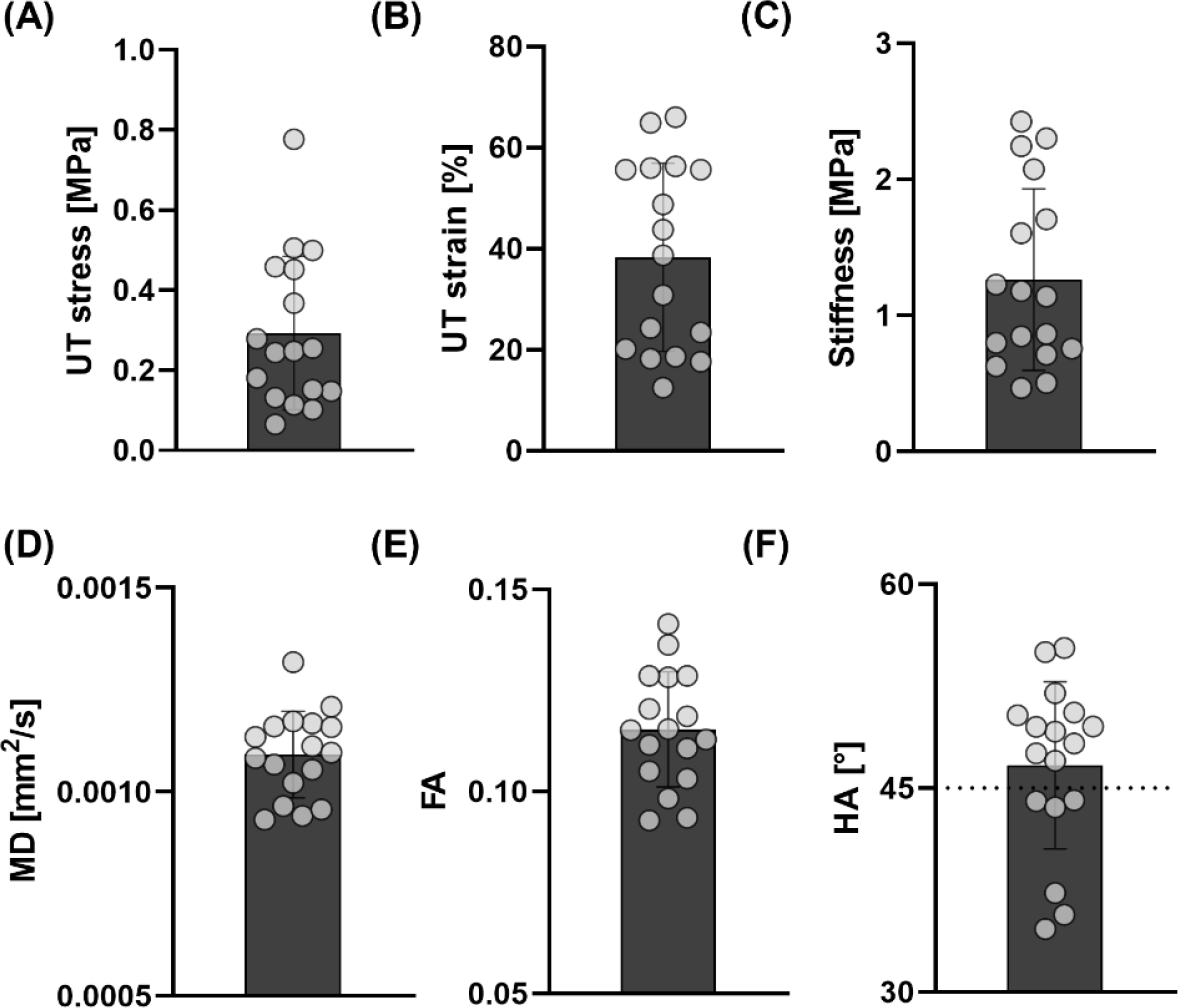
Mechanical properties and DTI-derived metrics of carotid atherosclerotic plaque strips. (A) UT stress, (B) UT strain and (C) stiffness of carotid atherosclerotic plaque strips and the corresponding DTI-derived (D) MD, (E) FA, and (F) HA. Dashed line in (F) is at 45°.

The UT strain calculated from the grip-to-grip separation was significantly higher than the measured DIC strain on the tissue surface, see Figure 4(A). However, the strains seen at the local failure locations on DIC were significantly higher than the mean strain across the tissue surface. When looking at how these strains correlate to the DTI-derived HA, the DIC gauge strain demonstrated the strongest correlation, Figure 4(C), followed by grip-to-grip strain, Figure 4(B), and then the DIC local failure strain, Figure 4(D).

**Figure 4.**
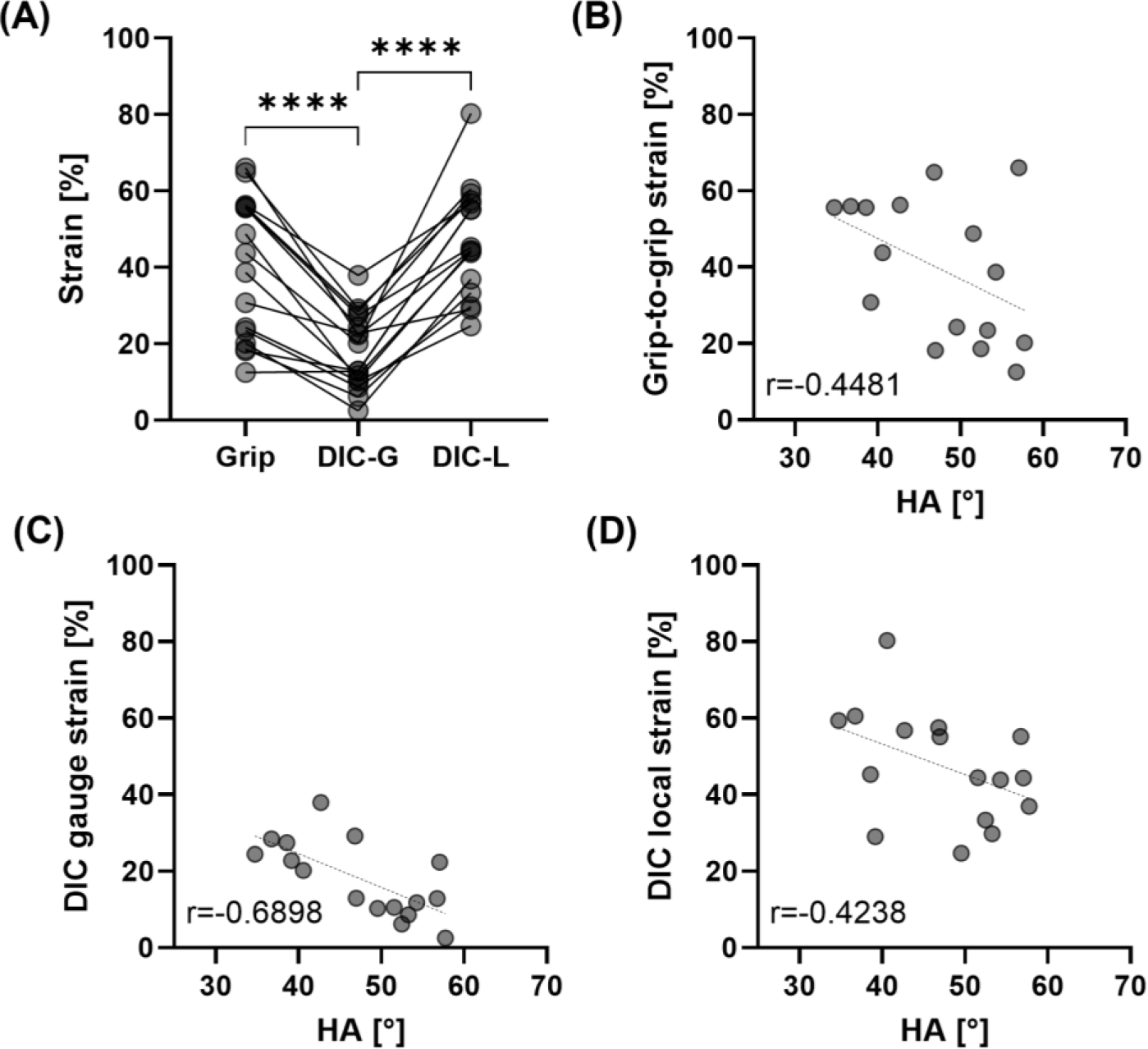
Strain measures and their correlations to DTI-derived HA. (A) Engineering strain from grip-to-grip (Grip), the gauge length from DIC (DIC-G), and local failure on DIC (DIC-L). Significance determined by repeated measures one-way ANOVA with Tukey’s post hoc multiple comparisons; ****p<0.0001. Correlations between DTI-derived HA and (B) grip-to-grip strain, (C) DIC gauge strain, and (D) DIC local strain.

Tractography of individual circumferentially cut strips highlighted the presence of four different microstructural alignments, see Figure 5. While most samples showed some circumferential alignment (Figure 5(1-3)), n=4 strips showed predominantly circumferential alignment with sparse axial tracts on the luminal side (Figure 5(1)) and n=4 strips were circumferentially aligned with the presence of a plaque cap shoulder (Figure 5(2)). Figure 5(3) shows n=5 strips which had circumferentially aligned tracts on the more medial side and a thick mixed alignment visible on the luminal edge, and n=4 strips with no clear alignment in Figure 5(4).

**Figure 5.**
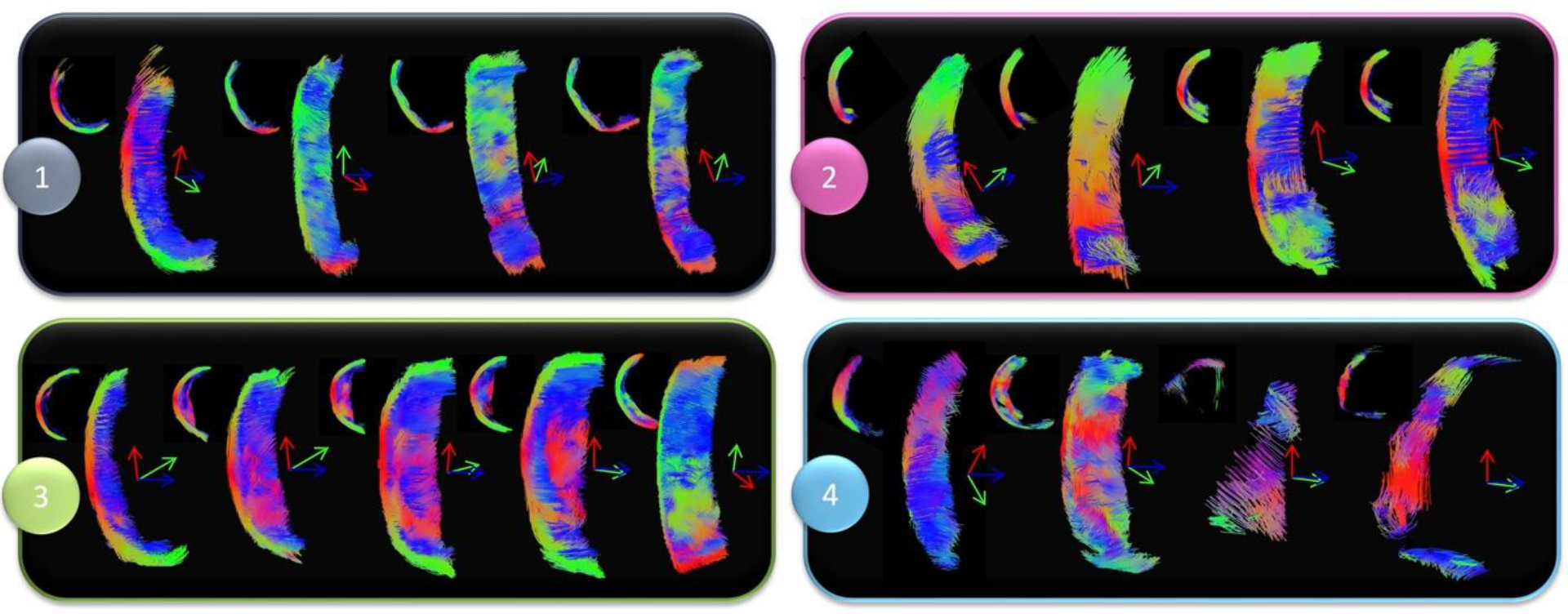
Tractography groupings of atherosclerotic plaque strips. (1) Predominantly circumferentially aligned tracts with sparce axial tracts on the luminal edge (concave side). (2) Predominantly circumferentially aligned tracts with the presence of a plaque cap shoulder – visible at the junction between fibres on the luminal side. (3) Circumferentially aligned medial tracts with large regions of mixed microstructural alignment on the luminal side and (4) overall mixed samples with no clear alignment. Red-green tracts indicate circumferential alignment, while blue is out-of-plane, axially aligned tracts.

Using the tractography based groupings shown in Figure 5, significant mechanical insight was uncovered, see Figure 6. The FA in mixed Group 4 strips (0.102 ± 0.01) was significantly lower than that in Group 3 (0.135 ± 0.008) and 2 (0.128 ± 0.009) strips, see Figure 6(B). The MD in Group 3 strips (0.0009 ± 0.00006 mm^2^/s) was significantly lower than that in Group 2 strips (0.0011 ± 0.00004 mm^2^/s). The HA was only significantly different between Group 2 and 4 strips, at 39.3 ± 5.32° and 53.3 ± 4.98°, respectively (Figure 6(C)).

**Figure 6.**
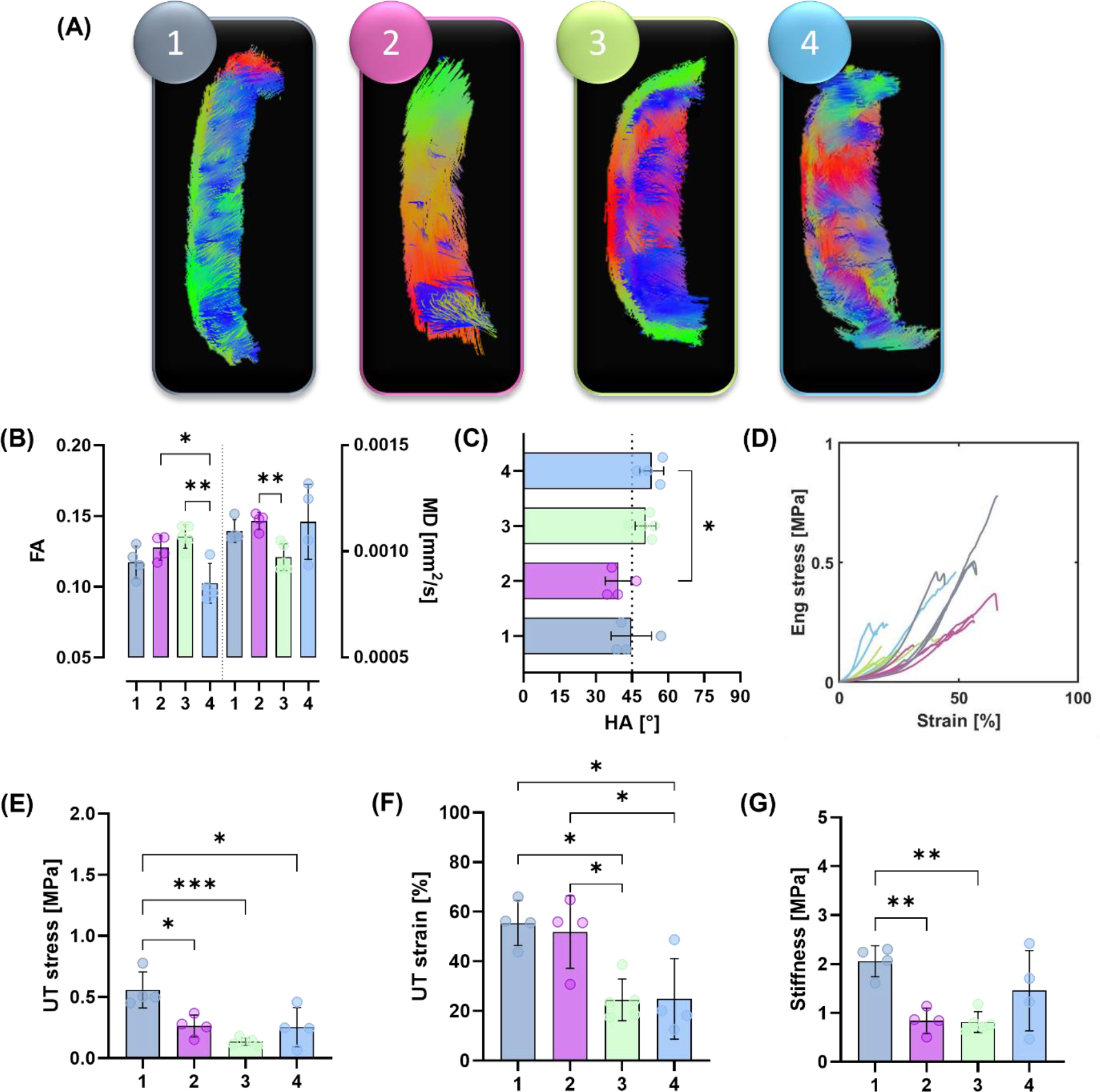
Mechanical properties of atherosclerotic plaque strips when informed by DTI-derived tractography. (A) Representative tractography of strip samples in each group. (B) FA and MD values within these groups; FA: significance determined by an ordinary one-way ANOVA with Tukey’s post hoc multiple comparisons, **p=0.0023, *p=0.0237. MD: significance determined by Brown-Forsythe and Welch ANOVA with Dunnett’s T3 post hoc multiple comparisons, **p=0.0089. (C) Mean HA for each group – the dotted line is at 45°; significance determined by ordinary one-way ANOVA with Tukey’s post hoc multiple comparisons, *p=0.0227. (D) Stress-strain curves for n=17 strips, colour coded by their respective groupings. (E) UT stress; significance determined by ordinary one-way ANOVA with Tukey’s post hoc multiple comparisons, Group 1 and 2 *p=0.0138, Group 1 and 3 ***p=0.0005, and Group 1 and 4 *p=0.0112. (F) UT strain; significance determined by an ordinary one-way ANOVA with Tukey’s post hoc multiple comparisons, Group 3 and 2 *p=0.0225, Group 3 and 1 *p=0.0145, Group 4 and 2 *p=0.0255, and Group 4 and 1 *p=0.0191. (G) Final stiffnesses; significance determined by Brown-Forsythe and Welch ANOVA with Dunnett’s T3 post hoc multiple comparisons, Group 1 and 2 **p=0.0051 and Group 1 and 3 **p=0.0053.

The stress-strain curves presented in Figure 6(D) illustrate the different mechanical behaviour between these strip specimen groupings. Predominantly circumferentially aligned strips in Group 1 have a significantly higher UT stress (0.559 ± 0.1 MPa), than those in Group 2 (0.264 ± 0.09 MPa), Group 3 (0.124 ± 0.02 MPa), and Group 4 (0.254 ± 0.2 MPa), see Figure 6(E). The UT strain (Figure 6(F)) of Group 3 strips (24.5 ± 8.43%), was significantly lower than that of Group 2 (51.8 ± 14.6%) and Group 1 (55.5 ± 9.11%) strips. Group 4 strips also had a significantly lower UT strain (24.9 ± 16.23%) than Groups 2 and 1. Figure 6(G) highlights the significantly stiffer response of Group 1 strips (2.06 ± 0.3 MPa) compared to both Group 2 (0.838 ± 0.2 MPa) and Group 3 (0.815 ± 0.2 MPa).

DIC not only allowed for strain contours to be displayed on the tissue surface but allowed for retrospective insight into how the strips failed. Figure 7 presents strain maps on representative strips for each group. High strain (shown in red) can be seen at the point of failure in Group 1 followed by abrupt failure (Figure 7(1)). Specimens in Group 2 consistently failed at the junction between differing microstructures present at the plaque cap before delaminating behind the lipid core (Figure 7(2)). Interestingly, the specimens in Group 3 similarly showed intimal tearing, however, they delaminated through the thickness of the mixed region (Figure 7(3)). Specimens in Group 4 failed quite variably. In the example strip shown, the location of failure did not show the highest local strain, showing that observable higher strains are not always co-located with failure.

**Figure 7.**
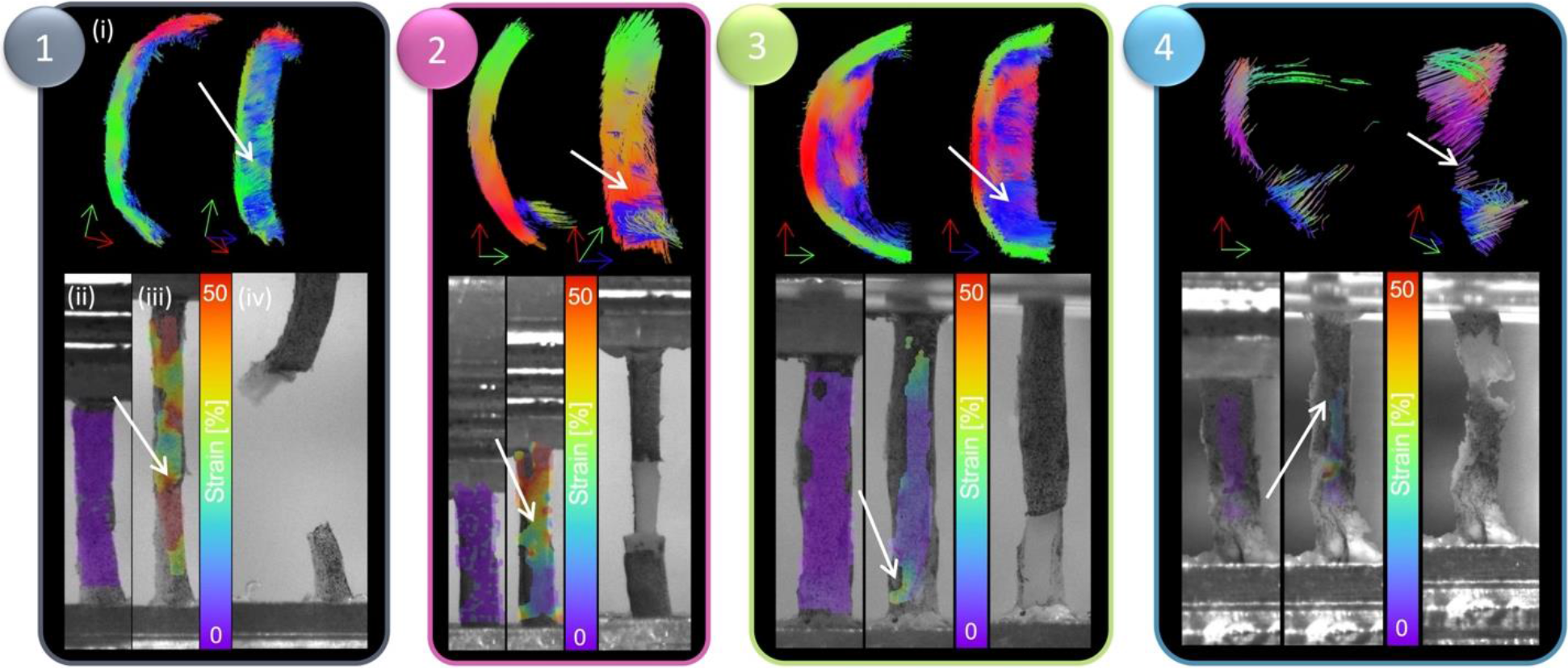
DIC strain contours and failure insights based on tractography groupings. For each grouping, (i) representative tractography is shown at the top, alongside strain contours on the DIC images at the (ii) reference frame (after preconditioning) and (iii) right before failure and a (iv) high-resolution image of the specimen right after failure. White arrows point to location of failure both on tractography and on strain maps.

Representative histology of the strips shown in Figure 7 are shown in Figure 8 to Figure 10. Figure 8 shows axial cross-sections for both strips in both Group 1 and 2. For both groups, failure occurred at the junction between differing microstructures, as pointed out by the green arrows. In the Group 1 sample, intimal thickening can be seen in H&E and decreased elastin is also visible in the Verhoeff’s-stained cross-section. PSR and PLM highlight the circumferential arrangement of this sample, as seen in tractography. Similarly, circumferential arrangement can be seen on the more medial side (left side of image) in the Group 2 sample. However, there is a distinct delineation of differing microstructure, where failure propagated behind the lipid core.

**Figure 8.**
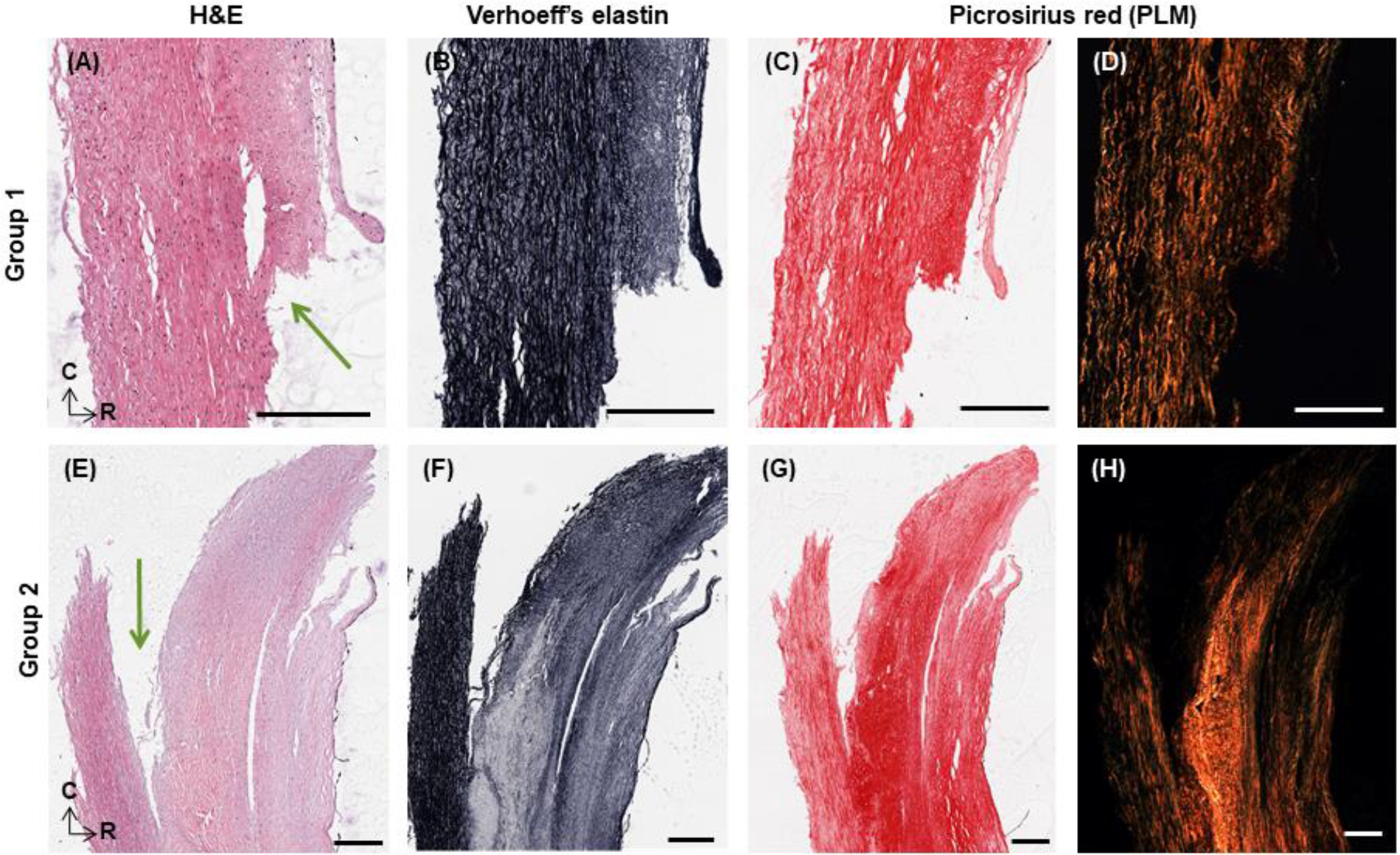
Representative histology of strips in (A-D) Group 1 and (E-H) 2. H&E, Verhoeff’s elastin, PSR and PLM are presented for each sample. Green arrows point to location of failure. Histology oriented to show axial cross-sections, moving from luminal edge towards media right to left and circumferential orientation top to bottom. All scale bars are 300 μm.

Figure 9 similarly shows representative histology for the Group 3 specimen shown in Figure 7. Unlike the cross-sections in Figure 8, these are radial slices through the thickness of the plaque wall due to difficulties orienting and embedding these mechanically tested strips. This specimen failed at the bottom edge of the strip, located at the bottom of the images, at what appears to be a plaque cap shoulder. The green box points to the centre of the strip (located in the centre of the gauge length) which is acellular, low in elastin, and has disorganised collagen (seen in PLM). The blue box highlights a plaque cap shoulder which could be like the shoulder at which the sample failed. Increased cell density is seen in this region, with nuclear alignment tending to be more longitudinally oriented. Distinct longitudinally aligned collagen fibres can be seen in PLM alongside regions of disorganisation.

**Figure 9.**
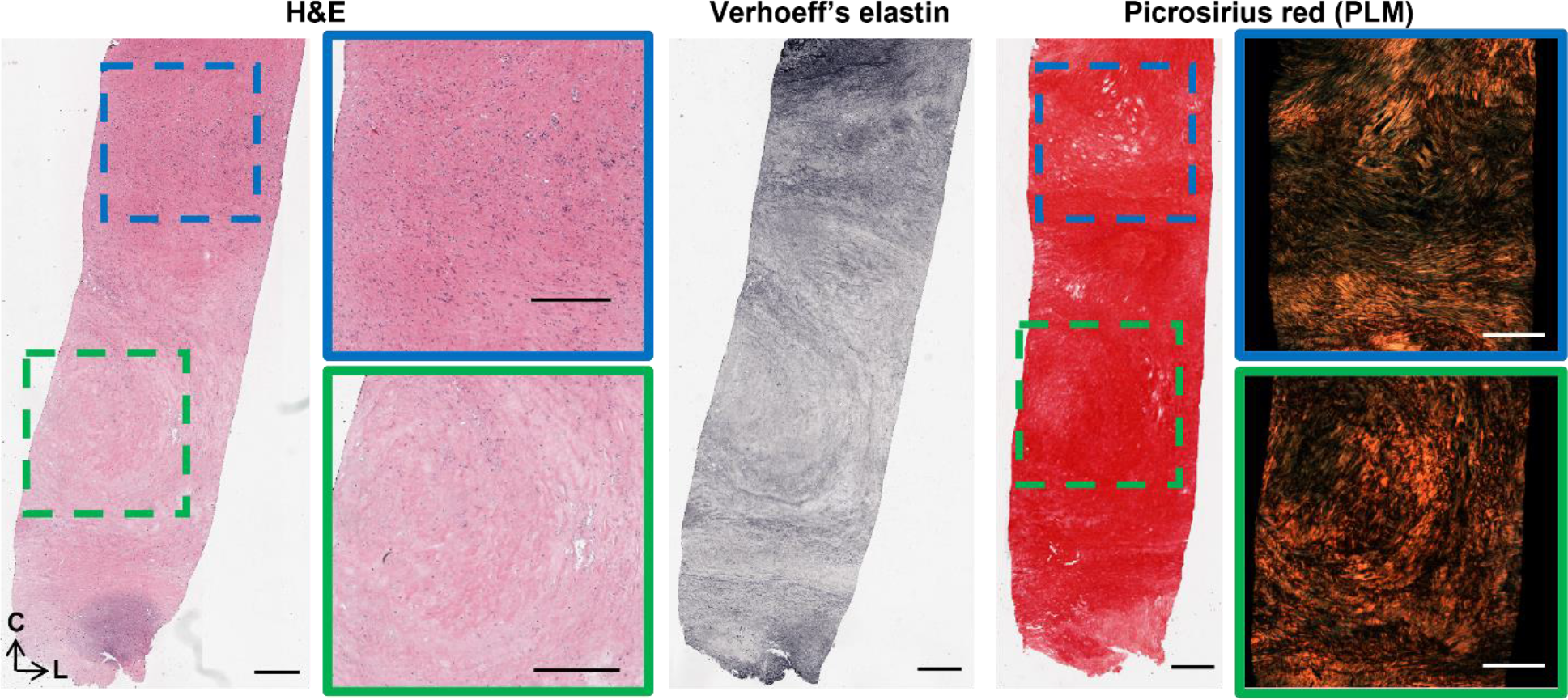
Representative histology of strips in Group 3. H&E, Verhoeff’s elastin, PSR and PLM are presented as radially sliced cross-sections: circumferential direction is top to bottom, while left to right is the longitudinal direction. Slices were taken at a depth approximately 140 μm from the luminal edge. Failure occurred at the bottom edge of the sample. All scale bars are 500 μm.

**Figure 10.**
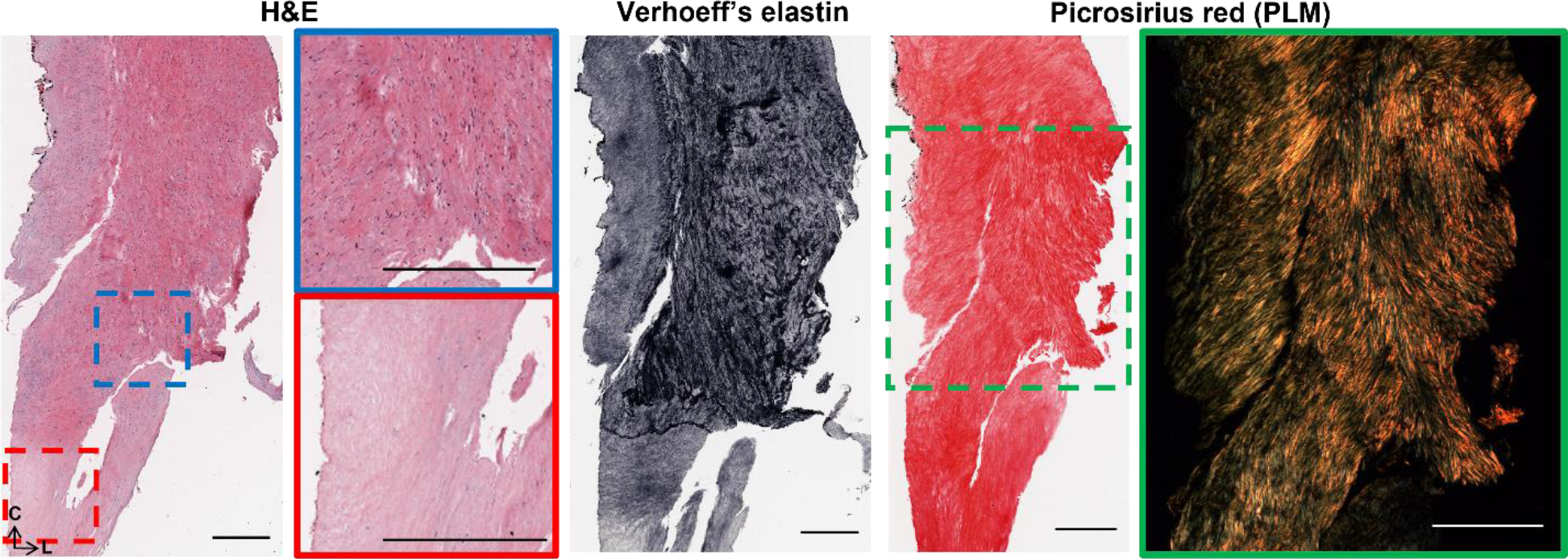
Representative histology of strips in Group 4. H&E, Verhoeff’s elastin, PSR and PLM are presented as radially sliced cross-sections: circumferential direction is top to bottom, while left to right is the longitudinal direction. Slices were taken at a depth approximately 140 μm from the luminal edge. Failure occurred from the blue box to the red box. All scale bars are 500 μm.

Lastly, Figure 10 highlights radial cross-sections of the Group 4 specimen seen in Figure 7. Failure delaminated down the length of the specimen, seen in Figure 7(4), and H&E presents the variable cell densities circumferentially. PLM shows collagen alignment tending towards circumferential but overall, quite disorganised.

## 4.0 Discussion

This study used DTI-derived metrics to investigate the mechanical integrity of carotid atherosclerotic plaques. Akyildiz et al. looked at tractography-derived fibre orientation in carotid plaques and, while variable, found the predominant direction to be circumferential^40^. Additionally, Opriessnig et al. used tractography to visualize the circumferential alignment both of the vessel wall and the plaque cap in one cadaveric carotid artery^49^. When assessing the overall tractography of plaques in this study, varying degrees of alignment can be seen (Figure 2). Variations in the microstructure can be seen both circumferentially, as well as longitudinally through the length of the plaque. These qualitative insights highlight the degree of disorganization and microstructural variation, not only between different plaques, but also within individual plaques.

The well-documented variable mechanical response of carotid plaques^20,21,24,26,50^ was also seen in this study. Comparatively, the UT stresses of circumferentially cut strips in this study are within the range of those determined previously^20,21,26^. More specifically, Groups 2, 3, and 4 failed at stresses lower than lightly calcified plaques^26,27^ and plaque caps with axially fibre alignment^25^. The strips in this study had no obvious evidence of calcifications and instead were fibrotic and lipid-rich, Figure 8(E-H) and Figures 9–10. Group 1 strips failed at significantly higher stresses, although not quite as high as plaque caps with circumferentially aligned fibres^25^ or the isolated medial layer of carotid plaques^47^. These strips were predominantly circumferentially aligned with respect to cells and collagen, although intimal thickening was evident (Figure 8(A-D)). While the stiffness observed in all groups in this study are higher than those reported for isolated plaque caps, they are within the ranges reported by Loree et al. of cellular atherosclerotic aortic tissue^17^. As evidenced in the tractography in Figure 5, the different groupings each demonstrated varying degrees of circumferential, axial, and mixed alignments. Interestingly, the Group 3 strips with thick mixed regions on the luminal side actually failed at lower stresses than moderately and heavily calcified plaques^26,27^. While calcifications have previously been pointed to as a vulnerable feature^51,52^, this finding highlights the significance of microstructural disorganization as well.

The strains for Group 1 and 2 strips are also in the range of previous studies^26,27,47^; however, Group 3 and 4 strips failed at lower strains more comparable to those of delaminated carotid plaque caps^25^. Previous studies pre-operatively used ultrasound to classify plaques as calcified, echolucent, or mixed and found no significance between groups. Only when using post-operative Fourier Transform Infrared analysis, was Mulvihill et al. able to differentiate between plaque compositions and their mechanical properties in pure shear tests^26^. Similarly, it was only when using tractography of individually tested atherosclerotic strips that different microstructures became apparent in this study (Figure 5). Ultimately, these microstructures yielded significant mechanical insight into the plaque tissue. The earliest sign of progressive atherosclerosis is the thickening of the intima, classified as an American Heart Association Type III lesion^53,54^. Group 1 strips in this study, defined as predominantly circumferential tracts with sparse axial diffusion on the luminal edge, showed signs of intimal thickening histologically (Figure 5(A) and Figure 8). These samples failed at significantly higher stresses and strains than the other microstructures in this study.

Following the thickening of the intima, AHA Type IV plaques or fibroatheromas can develop^55^. Thin cap fibroatheromas exhibit low smooth muscle cell density in the plaque cap which occludes a lipid or necrotic core^56^; however these atheromas often fail to narrow the vascular lumen despite thickening the arterial wall^57^. Both Groups 2 and 3 in this study exhibited the circumferential alignment known to be present in the medial layers of healthy arterial walls (Figure 5(B and C)), but also showed signs of more advanced plaque development. Group 2 strips were similar in microstructure to Group 1 but had the distinct presence of a plaque cap shoulder. Tractography was capable of visualising the plaque cap and also showed a circumferential alignment of the cap, see arrow in Figure 7(2), which was corroborated by the PLM histological images seen in Figure 8. Circumferential alignment is clear on both the luminal and medial edges, and surrounds an elastin-poor, disorganised (with respect to cell and collagen content) region. There also appears to be evidence of cholesterol crystals, which are believed to arise from cellular apoptosis^54^. Mechanically, these strips appeared to initially strain similar to Group 1 strips. When looking to the DIC images, the circumferentially aligned regions are bearing the load, until ultimately failure occurs at a significantly lower stress at the junction between the circumferential medial regions and the plaque cap shoulder (Figure 7(2)). Group 3 strips showed varying degrees of alignment in the plaque cap, but all show a thick, mixed region between the cap and the medial layers of the plaque. While both Groups 2 and 3 failed on the luminal edge of the plaque, Group 3 failed through the mixed region whereas Group 2 strips delaminated behind the lipid core – seen both in the DIC images and histologically (Figure 7(2 and 3) and Figure 8). Mechanically, Group 3 strips failed at significantly lower stresses and strains than both Groups 1 and 2 – identifying them as the most vulnerable microstructure of those seen in this study (Figure 6(E and F)). When looking histologically, it becomes clear that Group 3 strips also failed at the plaque cap shoulder – where there is a distinct difference in cell density and collagen orientation (Figure 9). The mixed microstructures in Group 4, unsurprisingly, show highly variable mechanical properties and a highly disordered microstructure via tractography (Figure 5(4)) and histology (Figure 10). Together, these results suggest that under the same physiological conditions, plaque tissue with a microstructural resembling that of Group 3 strips would be the most vulnerable to rupture. Conversely, a microstructural alignment mimicking that in Group 1 would be the least vulnerable and more stable – suggesting intervention may not be needed.

Another interesting finding from this work highlighted differences in the strain across the tissue. The strain calculated from the grip-to-grip separation was considerably higher than the strain across the tissue surface measured using DIC. However, the local failure strains on the tissue surface were significantly higher than these mean strains on the tissue surface. Previous work has shown that strain, rather than stress, might be a better indicator for plaque cap vulnerability^24,25^. The work presented here shows that the strain at local regions of plaque rupture might be significantly higher than the overall strain of the tissue.

While this study provides novel insights between a clinically relevant imaging technique and mechanical characteristics of human plaques, there are some limitations to the study. Firstly, sample numbers are limited. The imaging field of view confined usable strip specimens to a localised region and samples which could not be registered back to this field of view were excluded, as well as samples which failed near the grips. Despite these challenges, the limited sample numbers show promise given their significant differences. With increased plaque samples and individual strips, the groupings used in this study could be applied across entire plaques to determine the plaque’s vulnerability, rather than strips. DIC strain measures were not used after initial intimal tearing as the tissue surface with the speckle pattern was disconnected from the more medial layer which continued to bear load. Additionally, while the lengthy scan time allowed for in depth non-invasive characterisation, it will also lead to some tissue degradation. Despite this, all samples were subjected to the same scan times and their relative differences can be compared. Future use of echo planar imaging would speed up acquisition significantly and limit potential degradation. In the future it would be advantageous to utilise bi-axial testing or inflation testing of plaque tissue to gain more physiological insight into the mechanical response of the tissue. The incorporation of an MRI compatible bioreactor which allows for imaging before and after testing would yield novel insights into how the underlying microstructure is changing under loading and ultimately DTI-derived metrics, such as the helical angle, could be incorporated into finite element models^58,59^.

It would be naïve to not address the hurdle of clinical translation for a DTI sequence at the carotid bifurcation. Acquisition challenges stemming from cardiac and respiratory motion and scan duration mean translation is not a trivial task. However, this work provides the fundamental insights into DTI-derived metrics and the mechanical insight they can yield for future translational studies. Additionally, a number of in vivo studies have used diffusion weighted imaging to investigate plaque components^10,60–66^, already establishing the usefulness of measuring water diffusion within these tissues. Extending these acquisitions to incorporate directionality, while not trivial, has the potential to better ascertain plaque vulnerability due to the direct link to mechanical integrity.

## Conclusion

In this study, fresh human atherosclerotic plaques from endarterectomy surgeries were imaged ex vivo with a DTI sequence, mechanically tested, and investigated histologically. For the first time, this work identified a non-invasive MR imaging technique which could yield microstructural insight into atherosclerotic plaques and could ultimately be used to identify microstructures more at risk of rupture. These novel findings have the potential to drive continued research in non-invasive imaging techniques linked with mechanical characterisation to better identify plaque vulnerability.

## Acknowledgments

The authors would like the thank the Department of Vascular Surgery at St. James Hospital and at the Galway Clinic for supplying the human tissue used in this study. Specifically, Dr. Prakash Madhavan, Dr. Zenia Martin, Dr. Adrian O’Callaghan, Dr. Sean O’Neill, Dr. Lucian Iacob, Dr. Mike Bourke and Dr. Aoife Kiernan from St. James and Prof. Sherif Sultan and Dr Niamh Hynes from Galway Clinic.

## Author contributions

B.T. and R.D.J. collected all specimens from surgeries. B.T., A.J.S., and C.K. contributed to the development of the DTI protocol and B.T. and A.J.S. performed all MR imaging. B.T. and R.D.J. performed all mechanical testing. B.T. performed all data analysis, histological processing, and wrote the manuscript. C.L. conceived and supervised the study whilst C.L., B.T., R.D.J., A.J.S., and C.K. contributed to the study design. All authors reviewed the manuscript.

