## Supplementary material for "Microstructural and mechanical insight into atherosclerotic plaques– an ex vivo DTI study to better assess plaque vulnerability"

Affiliations:
[1]Trinity Centre for Biomedical Engineering, Trinity Biomedical Sciences Institute, Trinity College Dublin, Ireland
[2]Department of Mechanical, Manufacturing and Biomedical Engineering, School of Engineering, Trinity College Dublin, Ireland
[3]Department of Medical Physics and Clinical Engineering, St. Vincent's University Hospital, Dublin, Ireland
[4]Trinity College Institute of Neuroscience, Trinity College Dublin, Ireland
[5]Advanced Materials and Bioengineering Research Centre (AMBER), Royal College of Surgeons in Ireland and Trinity College Dublin, Ireland

### Supplementary data

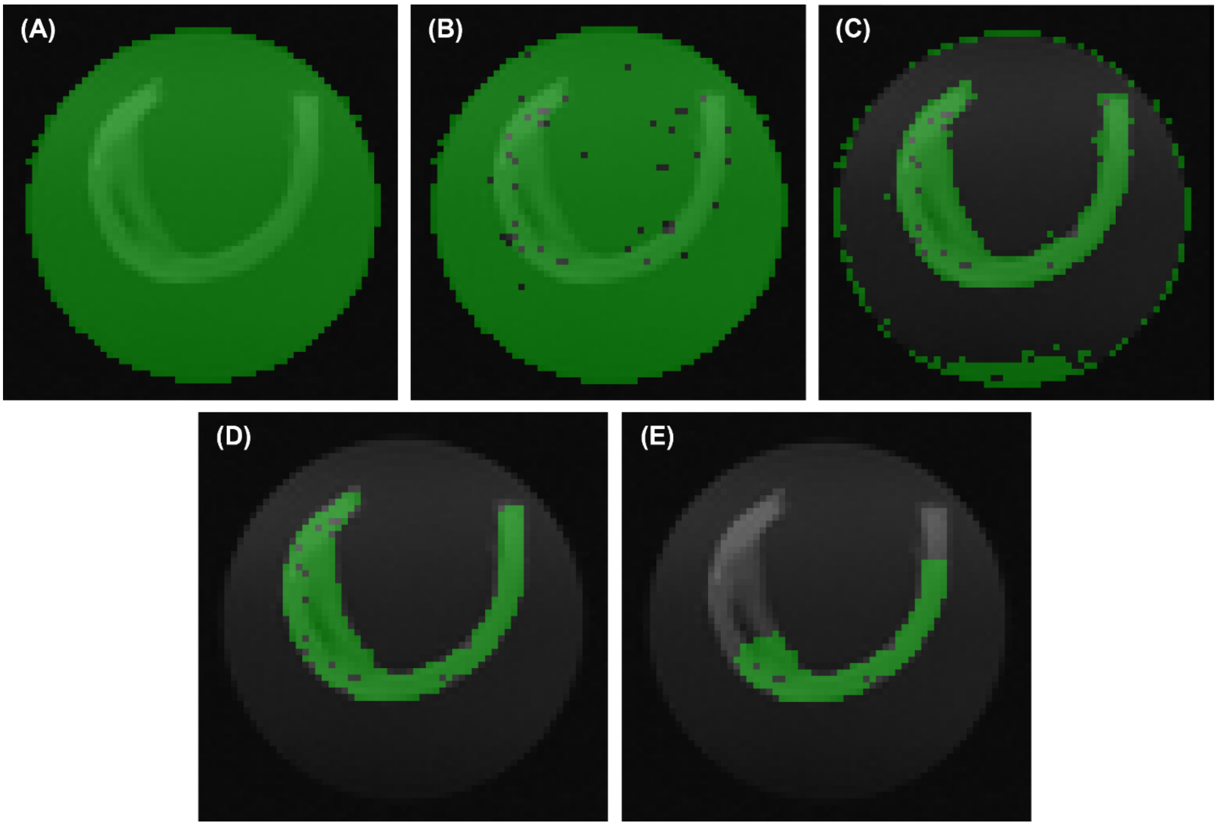


Supplementary figure 1. Masking removal of MR data. Green overlay is usable MR data mask after (A) low signal removal (removes signal outside falcon tube and calcifications), (B) high tensor residuals, (C) PBS removal, (D) manual stray pixel removal, and (E) removal of tissue within the grips during uniaxial extension.

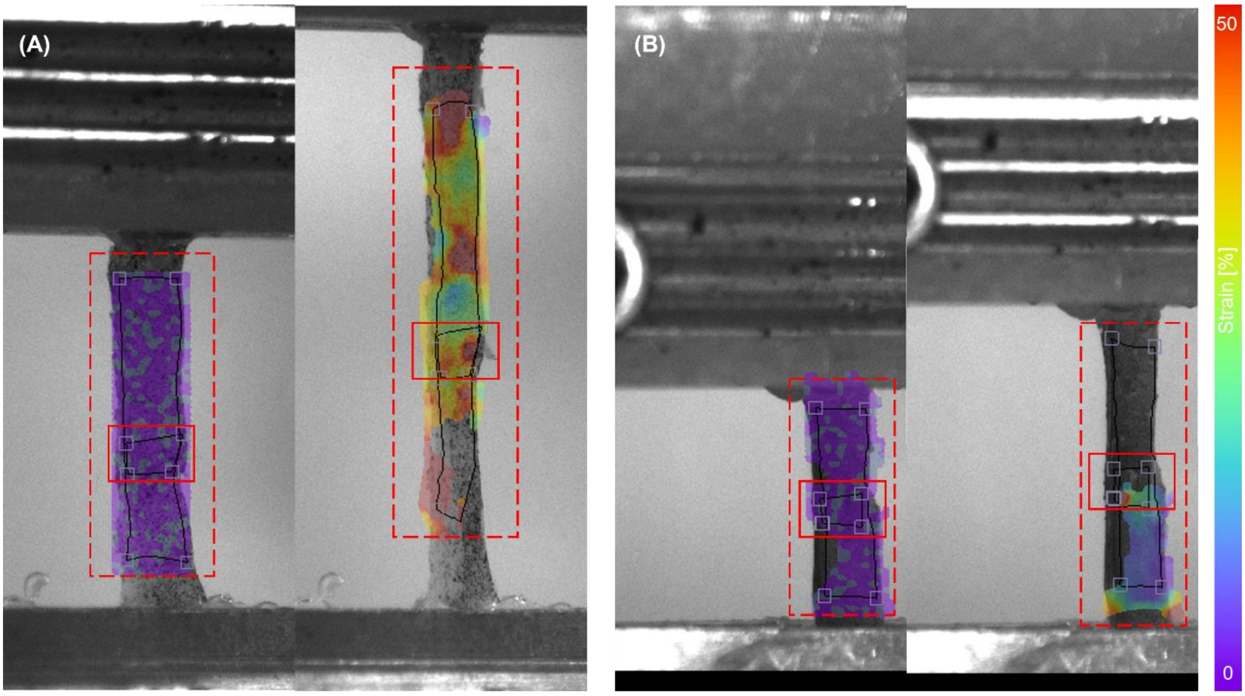


Supplementary figure 2. Solid lined boxes surround the region of interest for local strain measurements at the location of failure, while dashed lined boxes surround the region of interest for strain measurements across the gauge length. Two representative (A) and (B) samples shown.

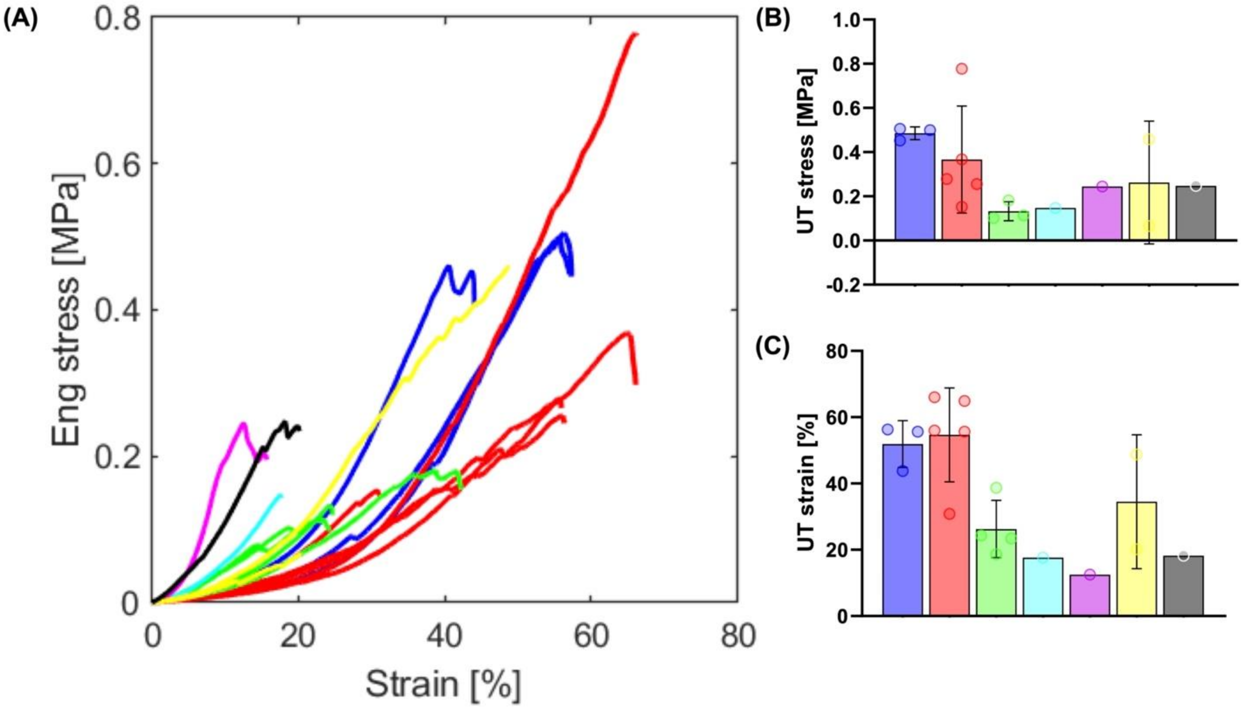


Supplementary figure 3. Mechanical properties of individual plaque specimens in this study. Colors in (A) the stress-strain curves correspond to the colors in the (B) UT stress and (C) UT strain graphs.
